# Wrong but useful: *Bombyx* silkworm W chromosome assemblies are flawed but still capture strongly reduced diversity of repetitive DNA

**DOI:** 10.64898/2026.07.21.739781

**Authors:** Martina Dalíková, James R. Walters

## Abstract

Degenerate sex chromosomes (*e.g.,* the Y or W) remain among the most difficult regions of eukaryotic genomes to assemble because they are highly repetitive and structurally complex. While many recent lepidopteran genome assemblies contain W chromosome scaffolds, the accuracy and consistency of these assemblies remain uncertain, due to lack of replication within species. However, the silkworm moth *Bombyx mori* is an exception, with numerous independent W chromosome assemblies currently available. We compared six independent long-read W chromosome assemblies, which proved to be highly inconsistent in structure, even among nominally identical genotypes. In contrast, autosomes and the Z chromosome were highly concordant among these assemblies, indicating that current assemblies remain unreliable for resolving W chromosome structure. Additionally, we analyzed repetitive DNA content across the genome. First, we combined assembly- and read-based repeat-discovery methods to generate a comprehensive and curated *Bombyx* repeat library, which we make publicly available. Assessing repeat content and diversity, we find that the W chromosome is comprised almost entirely of repetitive DNA but that the richness and divergence of W repeats are substantially reduced compared to the remainder of the genome. This reduced diversity, initially inferred from assemblies, is confirmed by direct analysis of PacBio HiFi sequencing reads partitioned by chromosome. We also demonstrate that the *B. mori* p50ma genome assembly (the current NCBI RefSeq assembly) carries a W chromosome and mitochondrial genome introgressed from *B. mandarina*. This discovery provided an opportunity to investigate patterns of divergence between closely related W haplotypes, revealing substantially more rapid turnover of repeat content on the W than elsewhere in the genome. Together, our results show that current W chromosome assemblies, although structurally flawed, nevertheless capture robust biological patterns of repeat diversity and support the hypothesis that rapid repeat turnover, rather than frequent chromosome replacement, may underlie the apparent lack of W chromosome homology across Lepidoptera.

## 1 Introduction

> “Remember that all models are wrong; the practical question is how wrong do they have to be to not be useful.”

in <u>Empirical Model-Building and Response Surfaces</u> (1987), by George E. P. Box & Norman R. Draper (p. 74)

Sex-limited chromosomes, such as the Y or W, are structurally and evolutionarily distinct, and often enigmatic in origin and function (Sahara et al., 2012; Bachtrog, 2013). These chromosomes are typically heterochromatic, gene-poor, and highly repetitive. Accordingly, they have remained recalcitrant to many DNA sequencing and assembly strategies that have supported myriad insights into the structure and evolution of autosomes and the homologous sex chromosomes (*e.g.,* X or Z) (Tomaszkiewicz et al., 2017; Chang and Larracuente, 2019; Rhie et al., 2023). Consequently, many outstanding questions persist regarding the patterns of diversity and underlying evolutionary processes that shape sex-limited chromosomes (Saunders and Muyle, 2024).

The W chromosome in Lepidoptera (moths and butterflies) embodies many of the analytical challenges and biological puzzles typical of degenerate, sex-limited chromosomes. Cytogenetic evidence clearly indicates that, in most species, the W is highly repetitive and heterochromatic, without detectable homology to the Z chromosome, and also that it evolves rapidly (Vítková et al., 2007; Sahara et al., 2012; Zrzavá et al., 2018). Indeed, the origin of the lepidopteran W is ambiguous and seemingly non-canonical, with the current emerging consensus that it reflects a co-opted supernumerary B chromosome rather than a degenerated homolog of the Z (Dalíková et al., 2017; Fraïsse et al., 2017; Voleníková et al., 2023). Early genomic studies of the W chromosome based on Sanger sequencing of a few BAC clones in the silkworm *Bombyx mori* supported the notion that it is overwhelmingly composed of repeats and largely devoid of protein-coding genes (Abe et al., 2005a).

In recent years, the advent of long-read sequencing has greatly increased capacity to meaningfully assemble and analyze degenerate sex chromosomes (Tomaszkiewicz et al., 2017; Chang and Larracuente, 2019; Rhie et al., 2023). In the case of Lepidoptera, this has yielded numerous new genome assemblies including many megabases of W-linked sequence (Lewis et al., 2021; Berner et al., 2023; Dai et al., 2024; Han et al., 2024; Lee et al., 2024; Orteu et al., 2024; Rueda-M et al., 2024; Wright et al., 2025; Shipova et al., 2026). These studies confirm the view that the W is replete with repetitive DNA, primarily transposons, and harbors very few, if any, protein-coding genes that are not transposons. This recent abundance of W assemblies has also spurred numerous efforts to investigate W chromosome homology between lepidopteran species, often in an effort to assess the hypothesis that the W is not widely conserved across species but rather reflects frequent independent origins (Lewis et al., 2021; Berner et al., 2023; Dai et al., 2024; Han et al., 2024; Orteu et al., 2024; Shipova et al., 2026). In a few cases where Z-W homology is evident, it does appear that a new W has arisen independently through a mechanism of “Single-Z Turnover” (Han et al., 2024; Shipova et al., 2026). Otherwise, the results of such comparative analyses are often ambiguous and sometimes contradictory, because W chromosome synteny and sequence similarity apparently erodes almost entirely even over moderate evolutionary distances, at least based on analyses using current assemblies, though this pattern is also reflected in cytogenetic studies (Vítková et al., 2007; Zrzavá et al., 2018). By contrast, autosomal synteny is readily detected between these same assemblies.

These observations raise critical questions about the evolutionary dynamics of the W chromosome as well as the quality of the assemblies on which these observations are founded. For instance, how quickly do W chromosomes diverge between species? Also, how accurately and consistently are we able to assemble the W chromosome in any given species? Furthermore, despite the availability of these W assemblies, little work has been done to characterize the diversity of repetitive DNA on the W chromosome relative to the remainder of the genome, limiting our understanding of W chromosome content, structure, and evolutionary dynamics. Where investigated, results indicate strikingly different patterns of repeat diversity on the W relative to the remainder of the genome, but the taxonomic breadth of these patterns is not well known, nor is the impact of any genome assembly artifacts on such inferences (Han et al., 2024; Shipova et al., 2026).

One straightforward way to establish some baseline answers to these questions is by examining independent assemblies of the W chromosome in the same or very closely related species. For such comparisons, it is reasonable to assume that W chromosome assemblies should be overwhelmingly consistent in structure and content. Radical differences between assemblies from the same species would highlight that strong technical artifacts may negatively influence comparative inferences. Also, differences in W chromosome content between recently diverged lineages can inform the tempo and mode in early stages of W chromosome divergence, assuming assemblies are sufficiently reliable to accurately represent such differences.

Despite the increasing availability of draft W assemblies, it remains rare to have multiple independent assemblies from a single species. However, one current notable exception is the silkworm, *Bombyx mori* (Table 1). The major economic significance of this species has placed it at the vanguard of lepidopteran genetics and genomics, motivating several recent independent long-read genome sequencing projects targeting females, thus yielding several draft W chromosome assemblies (Ma et al., 2019; Zhang et al., 2022; Han et al., 2024; Lee et al., 2024; Wan et al., 2025).

**Table 1.**
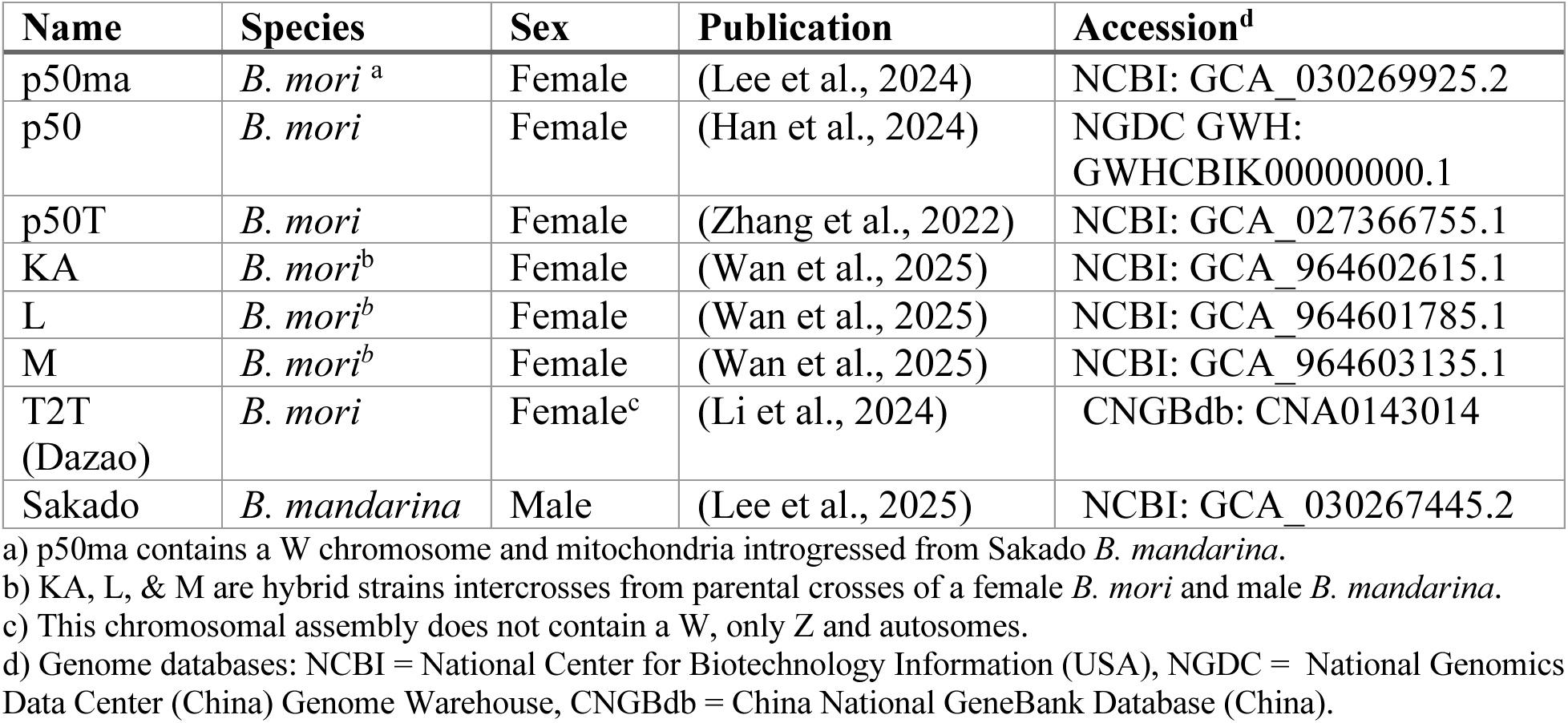
Bombyx genome assemblies employed in this research.

**Table 2.** Species-diagnostic W markers.

| Name | GenBank Accession | *B. mori* W | *B. mandarina* W | Target Length (bp) | Reference |
| --- | --- | --- | --- | --- | --- |
| Rikishi | AB126069.1 | Present | Absent | 959 | (Abe et al., 2005b) |
| W-Yamato | AB012908.1 | Absent | Present | 694 | (Abe et al., 2002) |
| Yukemuri-L | AB126065.1 | Short variant present | Short variant absent | 737 | (Abe et al., 2005b) |

These resources provide a novel opportunity to assess consistency in W chromosome assembly and to further characterize W chromosome content, which is the aim of our research presented here. Our analyses emphasize repetitive DNA content, because this is the outstanding feature of lepidopteran W chromosomes. Accordingly, we generated and make publicly available a comprehensive repeat database for *Bombyx*, integrating and consolidating repeats from both *B. mori* and *B. mandarina*, the “wild” silkmoth progenitor of *B. mori* (Goldsmith et al., 2005). To our knowledge, no such repeat database was previously available for widespread use, despite the prominence of *Bombyx* in genomics research.

Using this resource, we demonstrate that the *Bombyx* W chromosome possesses a profile of repetitive DNA substantially distinct from the rest of the genome. Additionally, we demonstrate that the current NCBI RefSeq *B. mori* genome assembly contains a W chromosome and mitochondrion introgressed from *Bombyx mandarina* (Lee et al., 2024). Finally, comparisons between *B. mori* and *B. mandarina* indicate differences in repeat content accumulate substantially more quickly on the W than elsewhere in the genome, consistent with the notion that rapid evolution of repeat content, rather than chromosomal turnover, may underlie the lack of sequence similarity frequently observed in comparative genomic analyses of lepidopteran W chromosomes.

## 2 Methods

### 2.1 Genome Assemblies

In total, our analyses employed data from eight different *Bombyx* genome assemblies, identified through literature and database searching (Table 1). We primarily focused on analyses of p50T, p50ma, T2T (*Dazao*), and Sakado, but also employed the others in various ways depending on our analytical goals. Summarizing strain histories briefly, the p50 strain and its close derivatives (p50ma, p50T, *Dazao*) reflect a long-established inbred genotype that is widely used in silkworm genetics (Goldsmith et al., 2005; International Silkworm Genome Consortium, 2008). The KA, L, and M strains are recently developed, originating with an initial cross between a *B. mori* female and a *B. mandarina* male, followed by backcrosses to females of distinct *B. mori* strains, and ∼30 generations of subsequent intercrosses (Wan et al., 2025). Thus, the K, A, and LM strains should all carry a *B. mori* W chromosome, though likely with different haplotypes, but otherwise have highly recombined chimeric genotypes mixing *B. mori* and *B. mandarina* genomes. The Sakado strain was isolated from a natural population of *B. mandarina* in 1982, subsequently isogenized via inbreeding, and is now widely used for comparisons and crosses with *B. mori* (Lee et al., 2025). For instance, the National BioResource Project (NBRP) Silkworm stock center has generated “chromosome replacement” lines, substituting each chromosome in the p50 genotype with a chromosome from Sakado (NBRP, 2024; Lee et al., 2025).

In our analyses comparing repetitive DNA on the W to the remainder of the genome, we limited our focus to data from p50T, p50ma, and Sakado. For the Z and autosomes, we used data from p50T and Sakado to compare between *B. mori* and *B. mandarina*. To compare between these taxa for the W, we used data from p50T and p50ma because the p50ma W chromosome is a *B. mandarina* haplotype (see Results); the Sakado is a male assembly and lacks a W. Of the available assemblies, we judge these three data sets to be the most comparable and informative because they employed PacBio HiFi sequencing technology and reflect well-established genotypes.

### 2.2 Female genome alignments

The six assemblies containing a W chromosome were aligned using minimap2 (v2.30) with default parameters (Li, 2018). Alignments were performed separately for the W chromosome and Z+autosomes. The p50T assembly was used as a common reference, with the other assemblies as the query. Resulting alignments were visualized as dotplots generated with the pafr R package (v0.0.2) (Winter, 2025).

### 2.3 W marker genotypes

To assess whether genomes carried a W haplotype from *B. mori* or *B. mandarina*, we searched the W chromosome assemblies for markers diagnostic of each species, as referenced in the NBRP newsletter (NBRP, 2024). Marker sequences were aligned to genome assemblies using minimap2 with the asm5 preset. Alignments covering less than 80% of the query sequence were discarded. Remaining alignment lengths were plotted for each assembly, and W haplotype was evaluated based on the detection of these markers. (*N.B.,* the Yukemuri-L marker PCR amplifies two products, ∼500 bp and ∼1200 bp. Only the shorter fragment is diagnostic of the *B. mori* W. The larger fragment, which amplifies in both sexes, contains an insertion according to our genome alignments (Abe et al., 2005b; NBRP, 2024).)

### 2.4 Mitochondrion analysis

We assessed the mitochondrial haplotypes associated with the p50ma and p50T assemblies by comparison to publicly available mitochondrial genome sequences from *B. mandarina* (n = 38) and *B. mori* (n = 1152) (Supplementary Table S1). The p50T and p50ma assemblies submitted to NCBI both lacked a mitochondrial scaffold, so we *de novo* assembled the mitochondrial genome from the associated PacBio sequencing reads (GenBank SRA accessions in Supplementary Table S2). Using minimap2 we aligned all reads to the *B. mori* reference mitochondrion assembly (GenBank: NC_002355.1) and selected reads which mapped to the reference with at least 80% of their length. The selected mitochondrial read sets were assembled using Canu (v2.2) in PacBio HiFi mode (Nurk et al., 2020). The resulting contigs contained multiple mitochondrion assemblies arranged tandemly, so the contigs were manually reduced to a single monomer corresponding to a single complete copy of the mitochondrial genome. This was performed separately for data from p50T and p50ma, resulting in a representative mitochondrial genome sequence for each.

These two assemblies were subsequently aligned with other available *Bombyx* mitochondrial genome sequences. To facilitate multiple alignment, each mitochondrial sequence was duplicated, oriented consistently, and trimmed to share a common start and end. All sequences were aligned using mafft (v7.525 with max 4 iterations) (Katoh and Standley, 2013). The alignment was processed with Gblocks (0.91b) to remove poorly aligned and divergent regions (Castresana, 2000).

To assess genetic relationships among *Bombyx* mitochondrial sequences, a principal component analysis (PCA) was performed on an allele-frequency matrix generated with the *tab* function from the adegenet R package (v2.1.11) (Jombart, 2008). PCA was conducted on the resulting matrix using the prcomp function with mean-centering and unit-variance scaling. The first two principal components were retained for visualization.

### 2.5 Repeat discovery and analysis

We generated a comprehensive *de novo* repeat database for *Bombyx* by combining results from a few separate data sources and methods for repeat discovery. First, we analyzed three *Bombyx* genome assemblies (p50ma, p50T, and Sakado) with the Earl Grey repeat discovery pipeline (v6.3.2, with default parameters) (Baril et al., 2024). Second, to better capture tandem repeats with monomers 50- 2000 bp long, we also analyzed these genomes using Tandem Repeats Finder (TRF; v4.09) and Tandem Repeats Analysis Program (TRAP; v1.1) (Benson, 1999; Sobreira et al., 2005). Third, to complement these assembly-based approaches, we directly analyzed Illumina resequencing data from two *B. mori* Dazao strain females (Genbank accessions: SRR5828088, SRR5828090) using RepeatExplorer2 (Galaxy Version 2.3.12.1) (Novak et al., 2013). To prepare the Illumina reads for the RepeatExplorer2 workflow, we removed the adaptors, quality-filtered, and trimmed reads to a uniform size of 95 bp using trimmomatic (v0.39), followed by removing overlapping read pairs and down-sampling to 9x10^5^ read pairs (Bolger et al., 2014). RepeatExplorer2 was run with default parameters using the Metazoan v3 library for repeat classification. Repeats found by TRF or RepeatExplorer2 estimated to comprise <0.01% of the genome were discarded from further analysis. Results from these distinct approaches to repeat discovery were consolidated into a single non-redundant repeat database “library” using CD-HIT (v4.8.1) following the 80-80-80 rule to define distinct repeat families (Li and Godzik, 2006; Wells and Feschotte, 2020). Manual inspection of the results revealed the combined library contained a single very large contig comprised of several tandem repeats of the same sequence, which we reduced to a single monomer.

### 2.6 Repeat masking and comparative analysis

We performed repeat masking on assemblies and also, separately, on sequencing reads (see below) using our custom library with RepeatMasker (v4.1.5) running the rmblast engine with a cutoff of 250 (Smit et al., 2015). To facilitate comparison between different partitions of the genome, RepeatMasker was run separately on the Autosomes, Z, and W, with identical parameters. Repeat landscapes were generated from repeat masking results processed by the RepeatMasker utility script calcDivergenceFromAlign.pl, parameterized to estimate divergences without modifying transition counts at CpG sites. Counts of repeats occurring uniquely or shared between *B. mori* and *B. mandarina* were visualized as UpSet plots using the R package ComplexUpset (V1.3.3) (Krassowski, 2020).

### 2.7 Sequencing read analysis

To confirm our repeat analysis results from genome assemblies, we performed similar analyses on unassembled HiFi sequencing reads partitioned by chromosome. HiFi sequencing reads from p50T, p50ma, and Sakado were each aligned to the unmasked T2T assembly using minimap2 with the “map-hifi” preset (GenBank SRA accessions in Supplementary Table S2) (Li, 2018). Reads mapping with alignments ≥80% of read length to the Z or autosomes were assigned accordingly. To identify W-linked sequences, noting that the T2T assembly doesn’t include a W chromosome, we selected reads whose longest primary alignment to the unmasked T2T assembly covered less than 50% of their length. This *under 50 (u50)* criterion reflects the rationale that since the W chromosome is almost entirely composed of repeats, many W-linked reads will align partially to repeats located elsewhere in the genome, but will have poor “global” alignments off the W.

As a complementary approach, we performed the same minimap2 alignment to the T2T assembly after masking using our *Bombyx* repeat library. Reads which did not map to the masked T2T assembly but do have at least 100 bp aligning to the unmasked genome were selected as W-linked. This *masked* criterion reflects the rationale that W-linked reads (which are comprised almost entirely of repeat sequence) should not map to the masked genome. However, reads which also do not align at all to the unmasked T2T genome are likely contamination and should be excluded.

For both the u50 and masked criteria, putative W-reads were subsequently screened for contamination using the GenomeFLTR webservice, filtering to remove reads with >10% similarity to the contamination database (Dotan et al., 2023). Ultimately, we used reads from p50T to represent *B. mori.* For *B. mandarina*, we used W-linked reads from p50ma, but reads representing the Z or autosomes were from the Sakado assembly. We assessed our success in partitioning W-linked reads by aligning W-markers to the selected reads using minimap2. We counted the number of reads per chromosome containing alignments ≥80% the marker length W-Yamato and Rikishi for *B. mandarina* or *B. mori*, respectively.

We subsequently performed repeat masking and generated TE landscapes for the collection of reads assigned to the Z and to the W. We did not perform repeat masking on autosomes, because it was impractical to run repeat masking on a full genome’s worth of raw HiFi sequencing reads.

### 2.8 Expression analysis

We assessed differences in gonadal expression of repeats between sexes using available RNA-seq data from three samples each of larval ovaries and testes (GenBank SRA: SRR34018379, SRR34018378, SRR34018377, SRR34018382, SRR34018393, SRR34018404). Read counts were generated using Kallisto (0.50.1) with a reference consisting of our custom *Bombyx* repeat library merged with CDS regions of 5671 single copy genes determined by running BUSCO (v6.0, using lepidoptera_odb12 database) on the T2T assembly (Simão et al., 2015; Bray et al., 2016). Expression differences were evaluated via DESeq2 (1.48.2) with testes as the reference tissue (Love et al., 2014). BUSCO genes were included to help improve estimation of normalization factors and dispersion parameters in the DE analysis, but were not the focus of differential expression analysis and were subsequently discarded for gene set enrichment analysis (GSEA). The DEseq2 Wald test *Stat* parameter was used to rank repeats for GSEA using the fgsea (v1.34.2) R package (Korotkevich et al., 2021). We tested for differential enrichment of each repeat type (*e.g.,* LINES, LTRs, DNA, *etc*.) in ovary or testes, though excluded Unknown repeats and the Penelope repeats (only 4 members). No boundary was used for the p-value estimation.

## 3 Results

### 3.1 Analysis of genome assemblies

### 3.1.1 W chromosome assemblies are highly discrepant

To examine consistency among W chromosome assemblies, we performed genome alignments using six *B. mori* genome assemblies with W chromosomes. We observed a great degree of variation in structure between the different W chromosome scaffolds (Fig. 1). In contrast, the Z and autosomes were highly consistent among the different assemblies (Supplementary Fig. S1). This strongly suggests that, even with the advantages provided to the assembly process by long-read sequencing and contemporary assembly methods, it remains quite challenging to consistently, and presumably accurately, reconstruct chromosomal-length scaffolds for the W chromosome.

**Figure 1.**
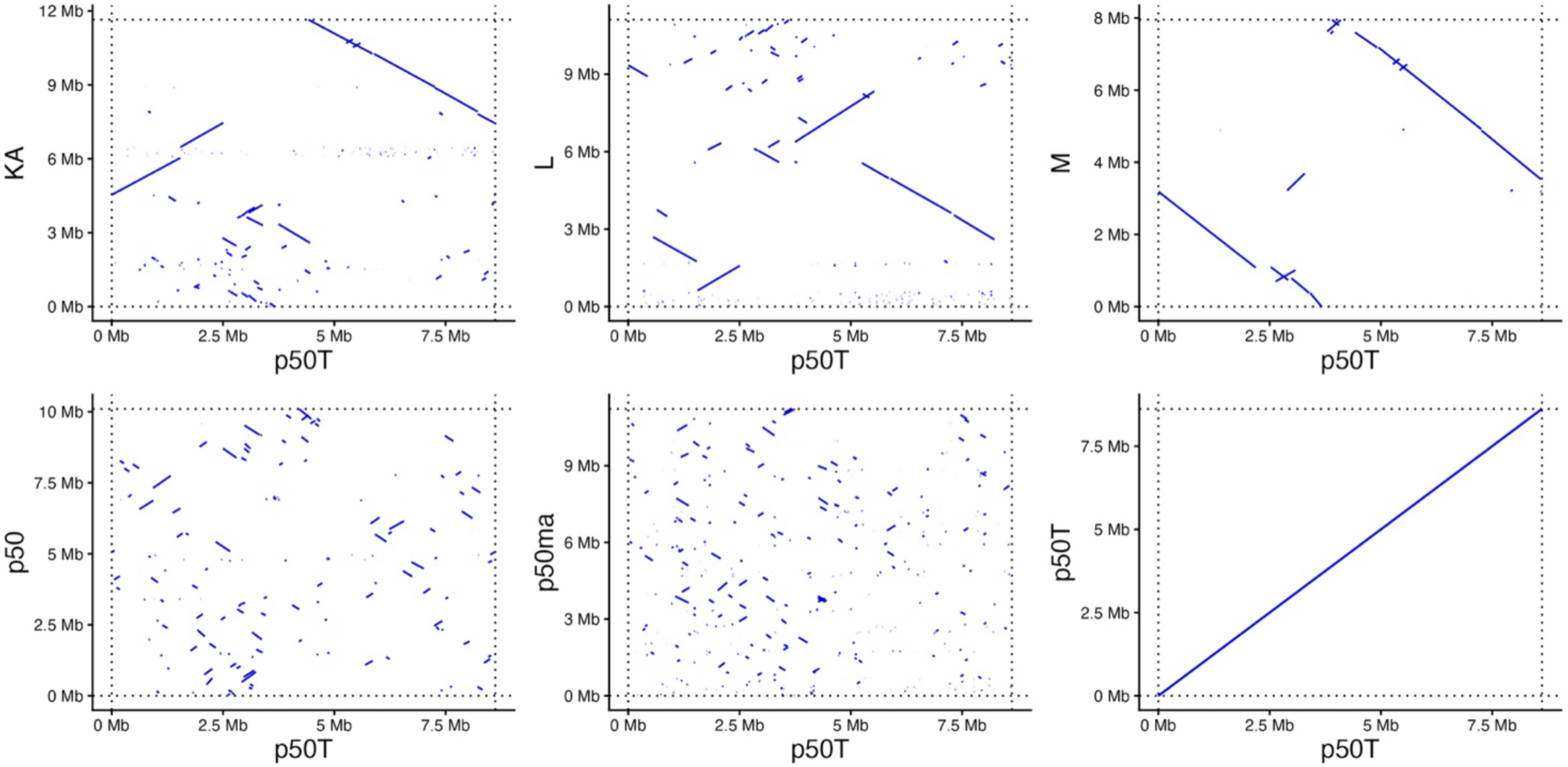
W chromosome alignments among different *Bombyx* genome assemblies. W chromosome scaffolds from various *Bombyx* genome assemblies were aligned using minimap2 to a common reference (p50T) and visualized as dotplots. The last alignment (bottom right) is of p50T with itself, provided as a comparison to the other alignments between assemblies.

It remains possible that the W chromosome differences among K, A, LM, and p50T assemblies reflect true biological variation, since these each have W chromosomes arising from different strains of *B. mori*. However, the naïve expectation is that p50, p50T, and p50ma should represent essentially identical genotypes and W chromosome haplotypes. Yet these three W assemblies are strikingly dissimilar. One straightforward explanation is that these discrepancies reflect artifacts of the assembly process, not biological variation. Nonetheless, we also explored in the literature what biological variation may distinguish these strains. While we maintain that mis-assembly is likely the best explanation here, we also uncovered that at least one of these, p50ma (the current NCBI RefSeq assembly), apparently carries a divergent W haplotype, which we sought to confirm.

#### 3.1.2 *B. mori* p50ma strain carries *B. mandarina* W and mitochondria

While investigating the histories for the p50-like strains, we encountered an important detail recently revealed about the history of the p50ma strain that seems not widely recognized among researchers outside Japan: this strain has been “contaminated” with a *B. mandarina* W chromosome. In an April 2024 Newsletter, the NBRP Silkworm stock center reported that the widely used *B. mori* p50T strain had been accidentally crossed with a chromosomal replacement strain carrying the W chromosome from the Sakado strain of *B. mandarina* (NBRP, 2024). The dispatch includes results from PCR-based markers and whole-genome resequencing that demonstrate the presence of the *B. mandarina* W, but that all other nuclear chromosomes appear derived from p50T. This contaminated strain carrying the *B. mandarina* W was subsequently named p50ma (Silkworm Base, 2026).

This history of contamination was not mentioned in the publication reporting the genome assembly for p50ma (Lee et al., 2024). Rather, the analysis and discussion presume the p50ma W sequence reflects a *B. mori* haplotype, leaving ambiguity about the exact nature of the W chromosome genotype present in this assembly. Accordingly, we performed our own assessment of the W genotype in the p50ma and other assemblies, searching for species-diagnostic W markers. Furthermore, we reasoned that p50ma should also carry a *B. mandarina* mitochondrial haplotype, since both the W and mitochondria are inherited maternally, and sought to assess this as well.

The NBRP newsletter reports two W-linked markers specific for *B. mori* (Rikishi and Yukemuri-L) and one specific for *B. mandarina* (W-Yamato), for which we searched in various *Bombyx* assemblies (Abe et al., 2002, 2005b). Consistent with p50ma carrying a *B. mandarina* W, we detected W-Yamato only in p50ma, while Rikishi and the “short” variant of Yukemuri-L were absent from p50ma (Fig. 2).

**Figure 2.**
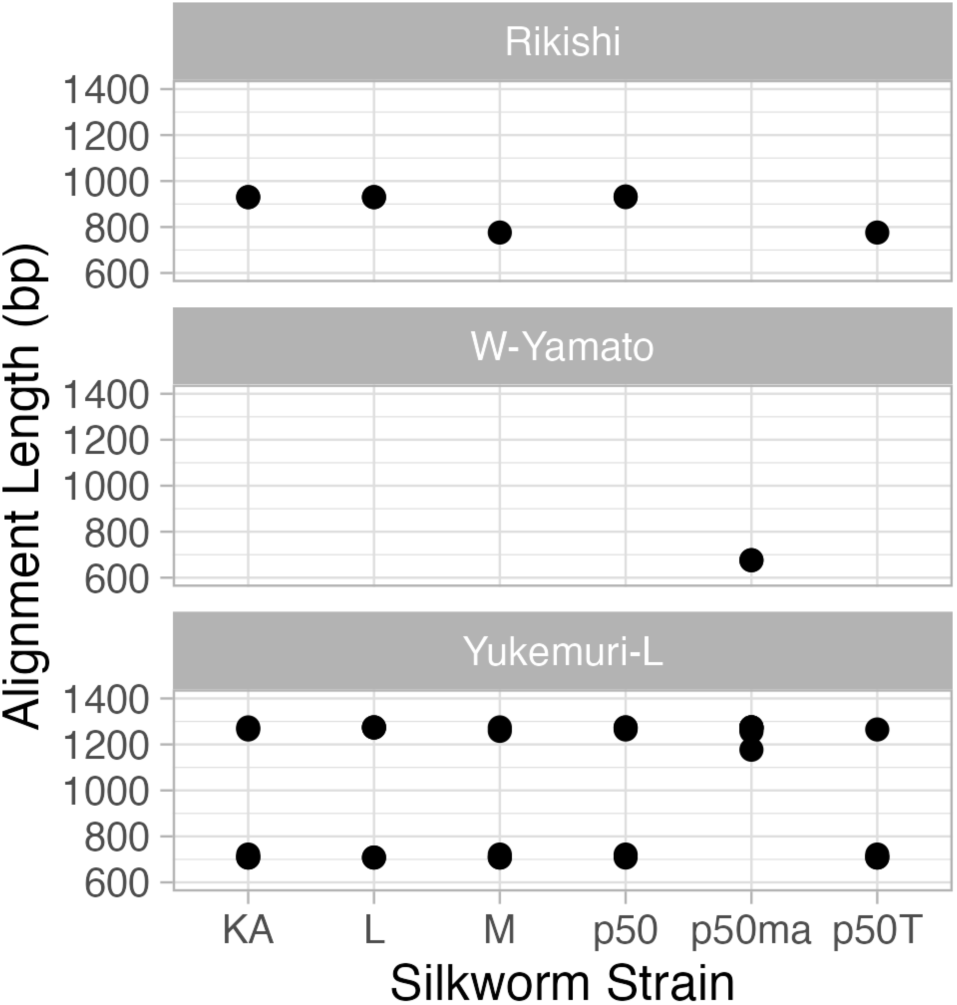
Detection of species-diagnostic W markers. Sequences of three W markers distinguishing *B. mori* from *B. mandarina* were aligned using minimap2 to the W chromosome of six *Bombyx* genome assemblies. The lengths of each individual alignment are plotted for each assembly. Alignments shorter than <80% of the query sequence were discarded.

Patterns of mitochondrial variation are also consistent with p50ma carrying a *B. mandarina* haplotype. To assess this, we first assembled mitochondrial genomes from the available sequencing data for p50ma and p50T. We then aligned these two assemblies with dozens of other *B. mandarina* and hundreds of *B. mori* publicly available mitochondrial genomes, analyzing the genetic variation with PCA. The PCA closely groups all strains of *B. mori*, including p50T, and indicates much greater diversity among *B. mandarina* mitochondrial genotypes (Chen et al., 2019). In the PCA, our p50ma mitochondrial assembly is quite distinct from the *B. mori* samples and clearly clusters with two other *B. mandarina* samples originating from Hokkaido and Tsukuba, Japan, respectively by PCA proximity (Fig 3).

**Figure 3.**
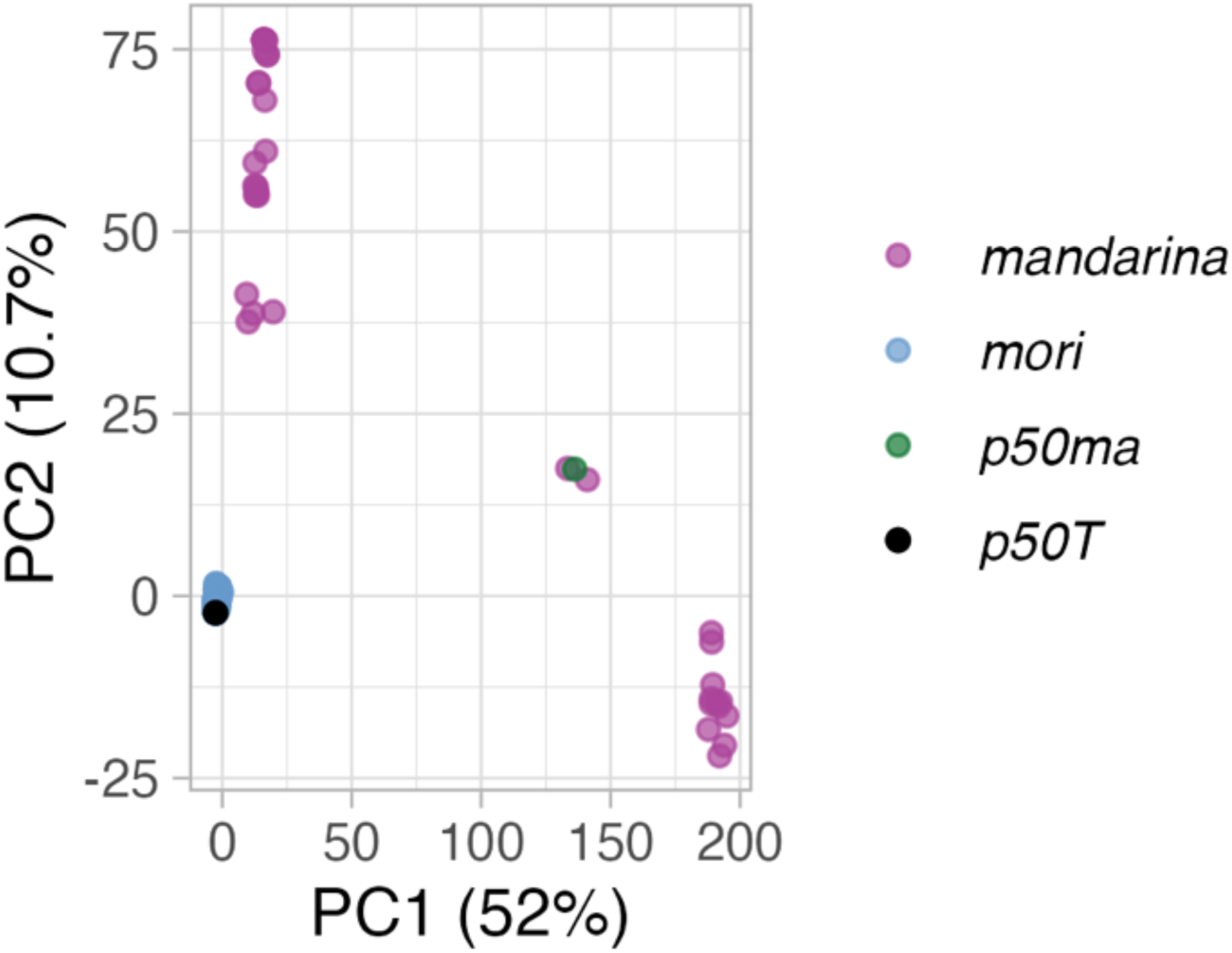
Principle Component Analysis of *Bombyx* mitochondrial diversity. Mitochondrial genomes were *de novo* assembled using PacBio HiFi reads from the p50T and p50ma assembly data. These were combined in a multiple alignment with publicly available mitochondrial genome sequences from *B. mandarina* (n = 38) and *B. mori* (n = 1152). Principle component analysis was performed on variation in the alignment summarized using the adegenet R package.

Based on these results, we presume that the W chromosome in p50ma is not a *B. mori* genotype and reflects a haplotype introgressed from the Sakado strain. In our subsequent analyses, we treat this W chromosome as representative of *B. mandarina*, in contrast to the *B. mori* haplotypes from other assemblies.

#### 3.1.3 A comprehensive *Bombyx* repeat database

One of our primary research aims was to characterize the patterns of diversity for repetitive DNA in *Bombyx*. While many recent publications reporting *Bombyx* genome assemblies perform repeat discovery and masking, we could not identify an easily accessible and publicly available relevant database of repetitive DNA. Furthermore, we needed our repeat analysis to capture repeats that might be specific to either *B. mori* or *B. mandarina*, which would not be the case with a library developed from only one of these taxa. To this end, we generated our own repeat library for *Bombyx*, merging repeat discovery approaches and data sets with the aim of comprehensively capturing repetitive DNA sequences, including both transposable elements (TEs) and tandem repeats (satellites). Our final *Bombyx* repeat library contained consensus sequences for 1778 distinct repeat families. Of these, 35% are Class I (RNA) TEs, 14% are Class II (DNA) TEs, 1% are satellites, and 50% remain uncharacterized (Supplementary Table S3).

This repeat library allowed us to proceed with comparing the repeat content of the W chromosome relative to the rest of the genome as well as between species. A RepeatMasker compliant FASTA formatted version of the library is available via Zenodo (see data availability statement) and the repeats were submitted for inclusion in the Dfam database (www.dfam.org).

#### 3.1.4 Disproportionate repeat content on the W chromosome

As is widely recognized for lepidopteran W chromosomes, the *Bombyx* W is comprised almost entirely of repeats (Sahara et al., 2012; Han et al., 2024). RepeatMasking with our *Bombyx* repeat library indicates >95% of the W chromosome assemblies correspond to repetitive DNA, which is about twice the amount of sequence masked elsewhere in the genome (Fig. 4a, Supplementary Table S4). There are also strong discrepancies in the proportions of repeat types found on the W versus the Z or autosomes (Fig. 4, Supplementary Table S5). For all chromosomes, retrotransposons (Class I TEs) strongly predominate. While LINEs show a proportional (e.g. two-fold) expansion on the W relative to other chromosomes, LTRs are strongly over-represented on the W, forming a proportion of masked sequence over six-fold greater than elsewhere in the genome. In contrast, SINES and DNA transposons (Class II TEs, including Helitrons) are strongly under-represented on the W relative to other chromosomes. Also on the W, ∼2-5% of masked sequence corresponds to satellites (e.g. tandem repeats) whereas satellite DNA is hardly detected anywhere else in the genome.

**Figure 4.**
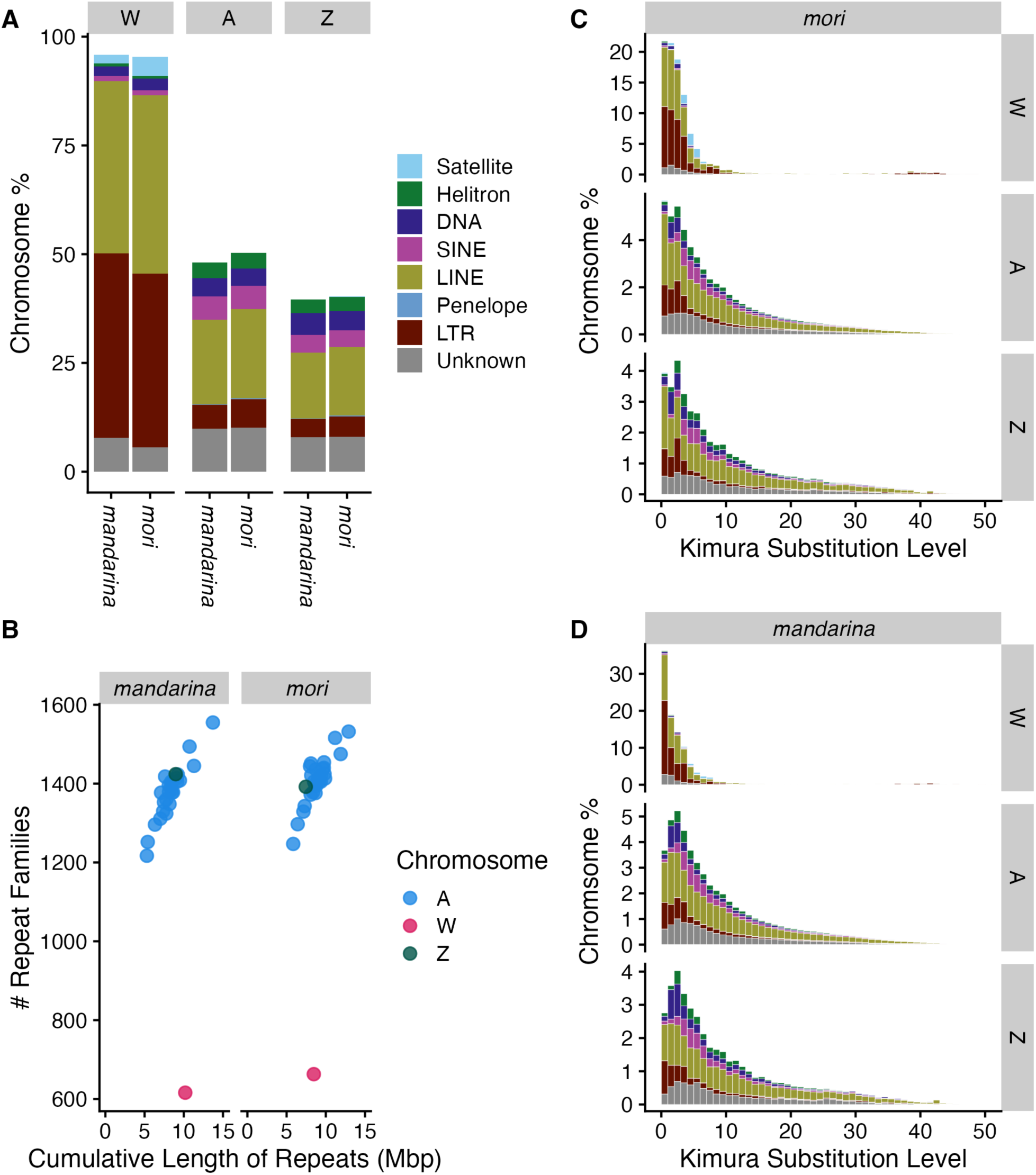
Patterns of repeat diversity in *Bombyx mori* and *B. mandarina*. (A) Proportions of repeat content for the W, Z, and autosomes partitioned by repeat type. (B) Relationship between repeat richness and the total amount of repeat content for each chromosome. (C) TE landscapes for the W, Z, and autosomes partitioned by repeat type for *B. mori* and (D) *B. mandarina*. Colors in TE landscapes (C,D) follow repeat types indicated in (A).

#### 3.1.5 Reduced repeat diversity on the W chromosome

We assessed repeat diversity on the W relative to the Z+autosomes by assessing two metrics: richness and divergence. Here we define *richness* simply as the number of different repeat families, analogous to the ecological concept of species richness (Moore, 2013). We assess divergence as the proportion of genetic differences between a specific TE genomic insertion and its corresponding consensus sequence, which can be interpreted as indicating the age of that TE insertion. The distribution of such divergences, often called a TE landscape, can highlight historical trends or other dynamics of TEs in the genome (Petersen et al., 2019). By both metrics, diversity of repeats on the W is strikingly reduced relative to the Z+autosomes.

For chromosomes other than the W, there is a clear, positive, approximately linear relationship between cumulative amount of repetitive DNA and the number repeat families on that chromosome (Fig. 4b). In other words, TE richness increases proportionally with total repeat content on the Z and autosomes. The W chromosome is a singular and striking exception to this pattern, with TE richness about half that of the other chromosomes, despite having total repetitive content comparable to other chromosomes. Thus, the W chromosome carries far fewer different repeat families than expected based on the remainder of the genome.

Furthermore, examining repeat divergence shows that TE insertions on the W are far more homogenous than elsewhere in the genome. We generated TE landscapes for representative *B. mori* and *B. mandarina* assemblies of the W, Z, and autosomes. The resulting TE landscapes are broadly similar between the two species. The most striking discrepancy is between the W versus the Z or autosomes (Fig. 4c,d). On the W, nearly all observed repeats show divergences <15% (though the W does contain a few repeats with divergences ranging up to 70%). By contrast, on the Z and autosomes, a substantial portion of the distribution extends up to ∼40% divergence.

This relative compression of the W chromosome TE landscape could indicate that TE copies on the W are substantially younger than elsewhere in the genome. However, it is also possible that some other molecular mechanism is homogenizing repeat content on the W, producing this pattern of reduced divergence. In any case, these results strongly indicate distinct processes are mediating the patterns of divergence among TE insertions on the W relative to all other chromosomes.

There are also some more subtle, but still noteworthy, differences between the two taxa. In *B. mandarina*, the Z and autosome landscapes peak at 3% divergence and decline sharply for less divergent repeats (Fig 4d). *B. mori* does not show this decline among the least divergent repeats, and for autosomes it has the greatest frequency of repeats in the least divergent tranche. This pattern suggests substantially more recent TE activity in *B. mori* than *B. mandarina*, which might possibly reflect reduced selection pressures on TEs or other dynamics associated with domestication.

However, this *B. mandarina* pattern of fewer “young” repeats on the Z and autosomes does not hold for the W chromosome. In fact, for both taxa, the least divergent tranche of W repeats has the greatest frequency. For *B. mandarina* in particular, this discrepancy in TE landscape profiles between the W versus Z or autosomes further highlights the idea that distinct evolutionary dynamics influence W-linked TEs.

#### 3.1.6 Differential expression of TEs between sexes

TEs are primarily expressed in gonads and are often differentially expressed in testes versus ovaries, a pattern that may associate with sex chromosomes (Chen et al., 2021; Warmuth et al., 2022). We explored this phenomenon in *B. mori* by analyzing TE differential expression in testes versus ovaries. Across all repeats, nearly half were differentially expressed, with roughly twice as many TEs over-expressed in ovaries than in testes (Table 3). GSEA indicated broad differences in expression of TE types between sexes (Fig. 5), with LTRs and LINEs significantly enriched (p < 0.004) in ovaries, but DNA elements significantly enriched (p = 0.02) in testes.

**Figure 5.**
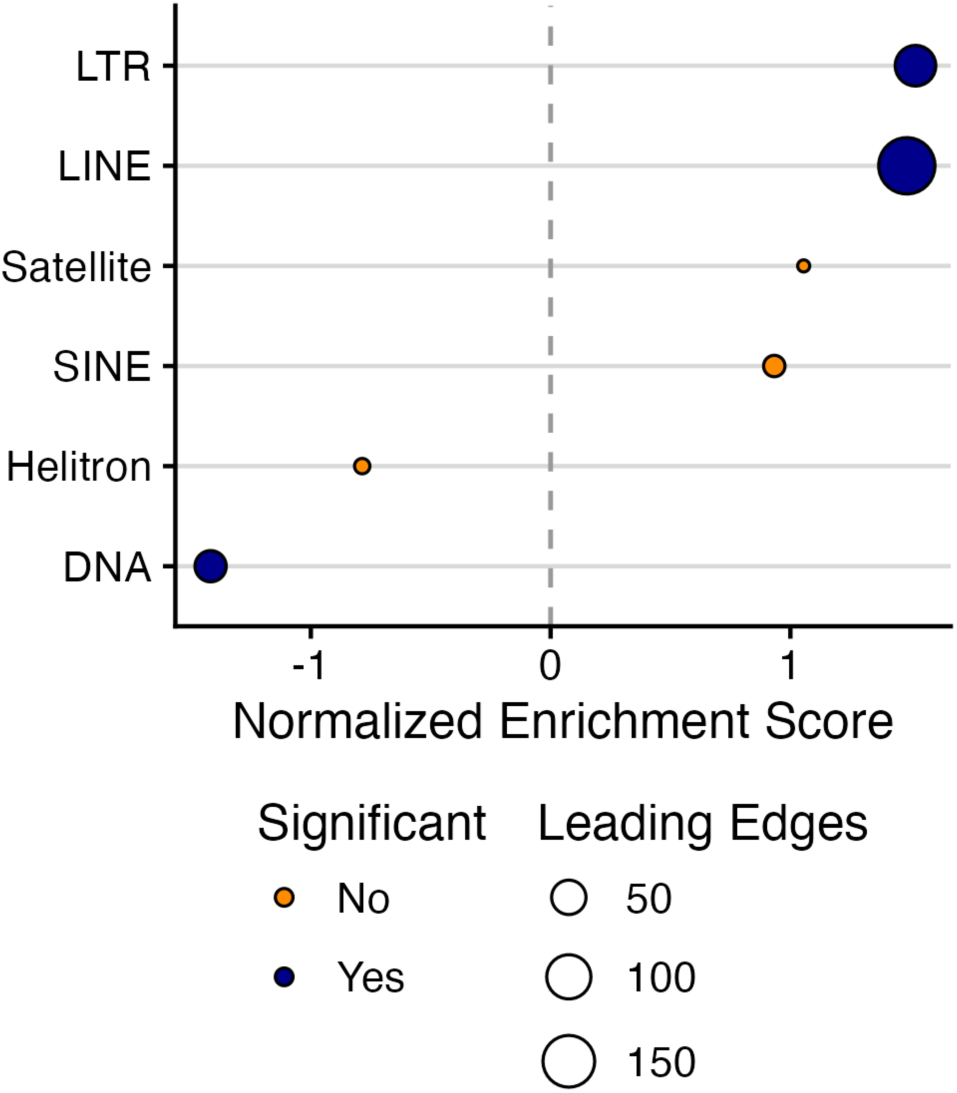
Gene set enrichment analysis (GSEA) results. Differential expression analysis of *B. mori* larval gonads in males versus females was used to perform GSEA on repeat families grouped by repeat type. A positive enrichment score indicates relatively greater expression in females. Statistical significance of enrichment reflects a Benjamini-Hochberg adjusted p-value < 0.05, indicated as blue coloring. Point size indicates the number of genes in the leading edge, the subset of genes contributing most strongly to the enrichment signal.

**Table 3.** Counts of significantly differentially expressed TEs upregulated in *B. mori* ovary or testes.

| Repeat Type | Ovary biased | Testes biased | Total in library |
| --- | --- | --- | --- |
| Satellite | 4 | 3 | 16 |
| Helitron | 21 | 9 | 83 |
| DNA | 53 | 40 | 173 |
| SINE | 11 | 1 | 32 |
| LINE | 153 | 34 | 378 |
| Penelope | 3 | 0 | 4 |
| LTR | 122 | 40 | 202 |
| Unknown | 220 | 115 | 890 |
| Total | 587 | 242 | 1778 |

These differences in TE expression, in conjunction with female-specific inheritance of the W, could play a role in shaping the relative abundances of TE types on the W versus the rest of the genome. As noted above, Class II (DNA) repeats are under-represented on the W, while LTRs are over-abundant (Fig. 4, Supplementary Table S4). Greater expression of DNA repeats in testes than ovaries, and vice versa for LTRs, would lead to the W being relatively less exposed to DNA repeats but over-exposed to LTRs. Over time, this pattern of exposure arising from the interaction of differential expression and female-specific inheritance could potentially produce the shifts in abundance of TE types occurring on the W relative to the remainder of the genome.

#### 3.1.7 Rapid evolution of repeat content on the W chromosome

Recognizing that the p50ma assembly captured a *B. mandarina* W haplotype presents the opportunity to investigate the relative rates of divergence on the W versus the other chromosomes at a very early stage of lineage divergence. Accordingly, we characterized each repeat family (as represented by distinct consensus sequences in our repeat library) as being shared or taxon-specific between *B. mori* versus *B. mandarina*. We observed that, as compared to the rest of the genome, the W chromosome has a vastly greater number of repeats detected in only one taxon (Fig. 5).

For autosomes together, the number of taxon-specific repeats is in the single digits, corresponding to less <1% of all detected repeats. For the Z chromosome alone, the number climbs to several dozen, or <10%. And for the W chromosome, it rises to over 100 repeats, or ∼36% in each taxon. However, when the entire genome is considered as a whole, the number of taxon-specific repeats is comparable to only the autosomes. This last observation contextualizes the result of observing many taxon-specific repeats when considering only the W chromosome. While W-linkage of these repeats has evolved quickly, the presence of these repeats in the genome is highly conserved. In other words, the W chromosome appears to gain and lose repeats at a much greater rate than other regions of the genome, but this reflects a kind of “dynamic equilibrium” relative to overall repeat content in the genome, at least at this very early stage of divergence.

### 3.2 Analysis of raw sequencing reads recapitulates assembly results

The large discrepancies we detected among the various W chromosome assemblies, which could potentially indicate substantial inaccuracies or incompleteness in W chromosome scaffolds, raises the concern that our assembly-based characterizations of repeat content on the *Bombyx* W chromosome may be flawed. Accordingly, we sought to recapitulate these results by directly analyzing the raw sequencing reads used in the assemblies, rather than the assemblies themselves. The large scale of data made it impractical to repeat mask sequencing reads from the entire genome, so we limited these read-based analyses to the Z and W only. For these two chromosomes, at least, the read-based analysis yielded results qualitatively similar to the analysis of assemblies presented above.

#### 3.2.1 Assignment of sequencing reads to chromosomes

We partitioned PacBio HiFi sequencing reads to chromosome by alignments to the T2T assembly. Reads aligning to the Z or autosomes were assigned accordingly, while reads which did not readily align to this male assembly were considered W-linked, following either the u50 or masked criteria (see Methods). Based on aligning W-specific markers to the resulting read sets, both approaches appear to have been effective at segregating W-linked reads. For both *B. mori* and *B. mandarina*, both criteria yielded at least several dozen reads assigned to the W containing the W markers, but the markers were never detected in reads assigned to any other chromosome (Supplementary Table S6).

#### 3.2.2 Read analysis yields reduced divergence of W repeats

Both criteria for selecting W-linked reads yielded TE landscapes highly consistent with results from analyzing the assemblies (Fig. 6). For both taxa, the distribution of W chromosome repeat divergences is almost entirely <15%, while repeats on reads from the Z reflect much greater divergences, ranging to near 30% or greater. Additionally, the read-based analysis also recapitulates the contrast between taxa in the least diverged tranche of repeats. In *B. mandarina*, the W is greatly enriched for these low-divergence repeat copies while the Z is depauperate in the same tranches. Meanwhile, in *B. mori*, the contrast is far less pronounced, with the frequencies in the least diverged tranche of repeats differing only slightly from the adjacent tranche.

**Figure 6.**
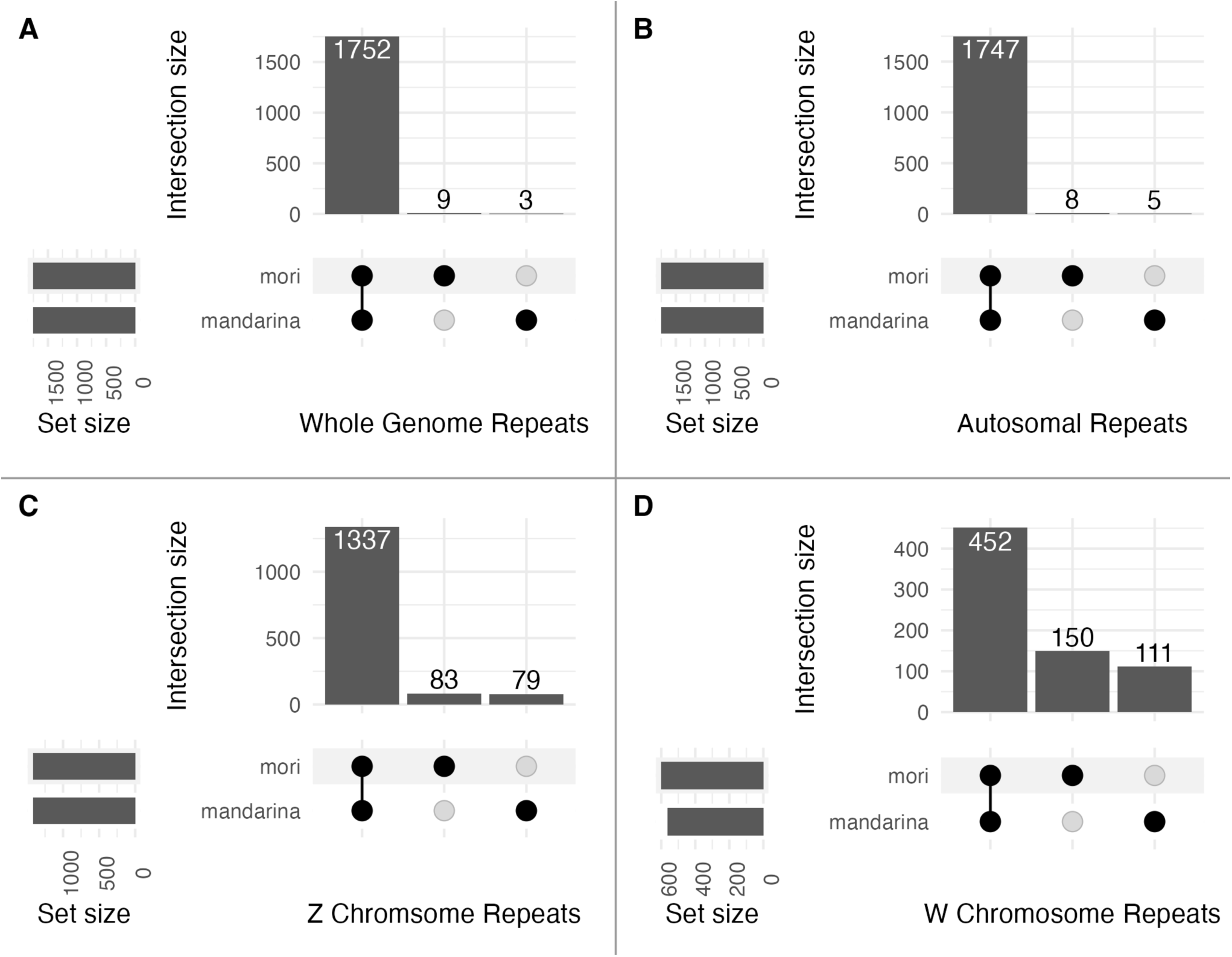
Shared and taxon-specific repeats detected between *Bombyx mori* and *B. mandarina*. (A) Whole genome (Autosomes, Z, and W). (B) Autosomes only. (C) Z chromosome only. (D) W chromosome only.

**Figure 7.**
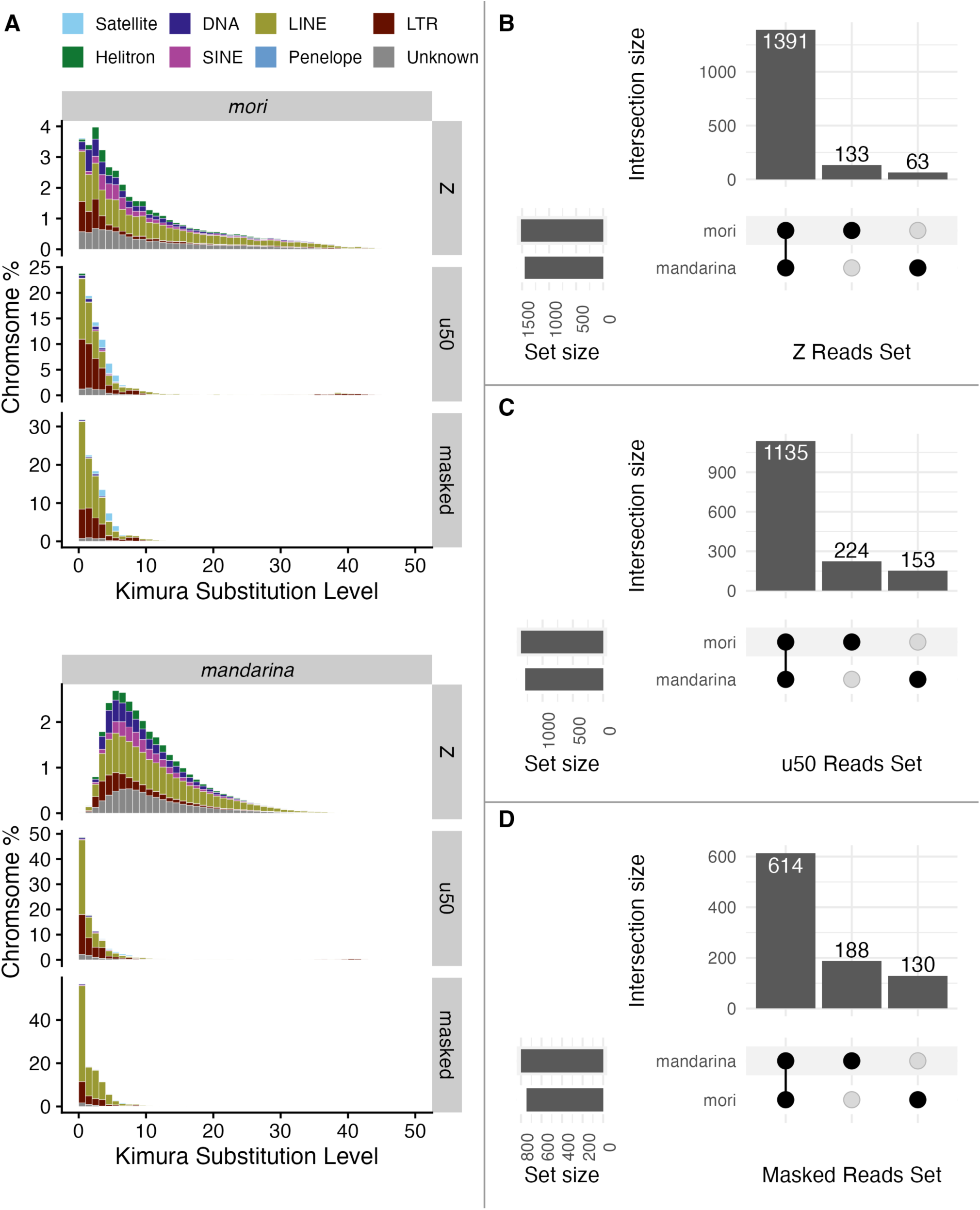
Read-based analysis of repeat diversity on sex chromosomes in *Bombyx mori* and *B. mandarina*. (A) TE landscapes for *B. mori* (top) and *B. mandarina* (bottom) produced for Z-linked reads and two different criteria for identifying W-linked reads. (B-D) Shared and taxon-specific repeats detected between *B. mori* and *B. mandarina* assessed from sequencing reads assigned to the Z (B) or the W using two different criteria for identifying W-linked reads (C,D).

#### 3.2.3 Read analysis indicates rapid evolution of W repeat content

Comparing shared versus taxon-specific repeats using read masking showed much greater specificity of repeats on the W than the Z. Taxon-specificity on the Z was 12%, about one half to one third that of the W, depending on the criteria for selecting W reads (*u50:* 24.9%, *masked*: 34.1%). These patterns are qualitatively similar results obtained from analysis of assemblies.

## 4 Discussion

The increasing availability of long-read genome assemblies is catalyzing opportunities to investigate the evolution, structure, and function of degenerate sex chromosomes (Tomaszkiewicz et al., 2017; Chang and Larracuente, 2019; Rhie et al., 2023). This is certainly true in Lepidoptera, where the enigma of the W chromosome’s history has motivated several comparative genomic analyses of the W chromosome that leverage the increasing abundance long-read assemblies (Lewis et al., 2021; Berner et al., 2023; Dai et al., 2024; Han et al., 2024; Orteu et al., 2024; Shipova et al., 2026). A recurring result from these analyses is that homologous sequence and structure is highly elusive between W chromosomes, sometimes prompting claims of frequent chromosomal turnover. However, such studies often do not sufficiently consider the possibility that the absence of detectable homology may in fact reflect extremely rapid sequence evolution (rather than chromosomal replacement and rapid degeneration), and also that substantial artifacts in assembly may be influencing these results.

Our results presented here highlight the importance of more fully considering these possibilities. First, comparisons between multiple independent W chromosome assemblies from closely related lineages or strains demonstrate that results can vary widely. While assemblies for Z and autosomes – “complex” and thus readily assembled regions of the genome – are highly reproducible and consistent, the repeat-rich W chromosome assemblies achieve at best only a few megabases of consistent assembly. These W assemblies probably are reliably capturing key features of its sequence content and composition (see below), but seemingly are still not consistently or accurately reproducing its overall structure.

Second, our analysis of repeat content demonstrates the potential for very rapid evolution of the W chromosome relative to the rest of the genome. Based on estimates for the domestication of *B. mori* from *B. mandarina*, these lineages have diverged for approximately 5000 years, a very modest amount of time in terms of genome evolution (Goldsmith et al., 2005; Chen et al., 2019). Yet it appears that >250 repeats are differentially present on the W chromosome – a difference several fold greater than the Z chromosome, which is otherwise expected (and observed) to be the most rapidly evolving chromosome in the genome (Meisel and Connallon, 2013; Sackton et al., 2014; Belleghem et al., 2018; Mongue et al., 2022). This elevated differentiation in repeat content was replicated in our analysis of sequencing reads, indicating it is a real biological pattern and not an artifact of assembly.

It remains unclear what processes may drive this rapid W evolution and otherwise shape patterns of diversity on the W chromosome. Female-specific inheritance interacting with differences in expression of TE types between ovaries and testes, as we observed, could be contributing towards differences in repeat types found on the W. While it is not clear what may initiate such differences in TE expression, the pattern would likely be self-reinforcing. For instance, if the W accumulates increased amounts of LTRs because of increased exposure to ovarian LTR activity, then the W itself may increasingly contribute towards that increased LTR activity.

Additionally, the lepidopteran W presumably shares characteristics typical of sex-specific chromosomes (*e.g.,* the Y) in other species, including suppressed recombination and a greatly reduced effective population size (Bachtrog, 2013; Sayres, 2018). These features alone could potentially result in increased rates of divergence. But the patterns of reduced richness and divergence we report here among W-linked repeats raise the potential of yet further complexity. Why should the W harbor a radically different proportion of repeat classes relative to the rest of the genome? And why are copies of these repeats so much more similar on the W than elsewhere in the genome, as prominently demonstrated in the *B. mandarina* TE landscapes?

This very low divergence among repeat copies on the W is not unique to *Bombyx*, as a similar pattern was also recently reported for three Lasiocampid moths (which are sister to Bombycoidea moths) and the much more divergent nymphalid butterfly *Hipparchia semele* (Kawahara et al., 2019; Shipova et al., 2026). In the case of the Lasiocampids, each species appears to harbor an independently evolved W, separately arising from Z-turnover. (Despite our general caution about invoking frequent W turnovers to explain a lack of homology, in this case, the phylogenetic analysis of Z-W gametologs reasonably supports the claim. Z-W homology in *H. semele* is much less clear.) It is striking that relatively young (the Lasiocampids) and, presumably, older (*Bombyx* and a butterfly) W chromosomes all share this pattern, though with distinct classes of TEs predominating in each taxon. For newly evolved W chromosomes, it may well be that the low divergence reflects many recent transposition events (Shipova et al., 2026). But this explanation is harder to reconcile with older W chromosomes where there has been greater time for mutations to accumulate and differentiate distinct TE insertions. If this pattern continues to be observed on lepidopteran W chromosomes, it would seem to require some further explanation beyond the TEs simply being young. One speculative possibility is that some form of concerted evolution is homogenizing the repeat sequences on the W (Elder, and Turner, 1995). Clearly there is opportunity for much future research addressing these unusual and unexpected patterns.

Of course, any future research addressing the lepidopteran W chromosome would benefit substantially from having highly accurate assemblies. There is clear precedent that this can be achieved through critical application of current technologies (*e.g.,* by combining ultra-long Nanopore with high-accuracy PacBio HiFi sequencing), so we are optimistic that such assemblies will be available soon for Lepidoptera (Rhie et al., 2023). Nonetheless, our results suggest that existing W chromosome assemblies are still highly informative for assessing many important aspects of chromosomal content and diversity. At least this appears to be true for analyzing repeats, which comprise the vast majority of the W chromosome, as evidenced by our qualitatively replicating from sequencing reads the key findings from our repeat analysis of assemblies.

Consequently, we want to emphasize the potential for rapid evolution of repeat content on the W chromosome to greatly obscure any signals of homology, even at moderate evolutionary differences where homology and synteny are readily evident elsewhere in lepidopteran genomes. Regarding W chromosome homology, it may be worth dwelling on the notion that “*absence of evidence is not evidence of absence*”. Rather than invoke rampant W turnover across the Lepidoptera to explain the frequent lack of clear W homology between species, we propose it is more likely the case that this typically represents a “Ship of Theseus” phenomenon (Gallois and Kurtsal, 2026). As in the Greek legend where a single ship is continually reconstituted with new planks as old ones rot away, we envision a scenario where the repeat content of the W chromosome changes so rapidly and thoroughly that it quickly becomes unrecognizable between species. Nevertheless, in this scenario, we would consider the W chromosomes homologous despite being comprised of entirely new sequences.

Whatever underlies the distinct evolution of lepidopteran W chromosomes, certainly increasingly accurate assemblies will be helpful to clarify the processes. In the meantime, the draft assemblies currently available may be imperfect, but nonetheless appear to be useful in accurately capturing prominent features of W chromosome diversity and function.

## Supporting information

Supplemental Table 1: Mitochondrial accessions

## 5 Conflict of Interest

*The authors declare that the research was conducted in the absence of any commercial or financial relationships that could be construed as a potential conflict of interest*.

## 6 Author Contributions

Martina Dalíková: Conceptualization; Data Curation; Formal Analysis; Funding Acquisition; Investigation; Methodology; Software; Visualization; Writing – Original Draft Preparation; Writing – Review & Editing.

James R. Walters: Conceptualization; Data Curation; Formal Analysis; Funding Acquisition; Investigation; Methodology; Project Administration; Resources; Software; Supervision; Visualization; Writing – Original Draft Preparation; Writing – Review & Editing.

## 7 Funding

MD was supported by the Kansas University Center for Genomics Postdoctoral Fellowship.

## 8 Acknowledgments

We are grateful to the organizers and colleagues at the 2025 12th International Workshop on Molecular Biology and Genetics of the Lepidoptera for constructive and informative conversation regarding this research.

## 9 Data Availability Statement

The *Bombyx* Repeat Library generated for this study can be found at https://doi.org/10.5281/zenodo.21361476.

The mitochondrial genome assemblies generated for this study were deposited in GenBank and be found under accessions XXXXX (p50T) and XXXXX (p50ma).

Scripts for data analysis and visualization are available via github: https://github.com/WaltersLab/Bombyx_W_analyses

## 1 Supplementary Material

**Supplementary Figure S1.**
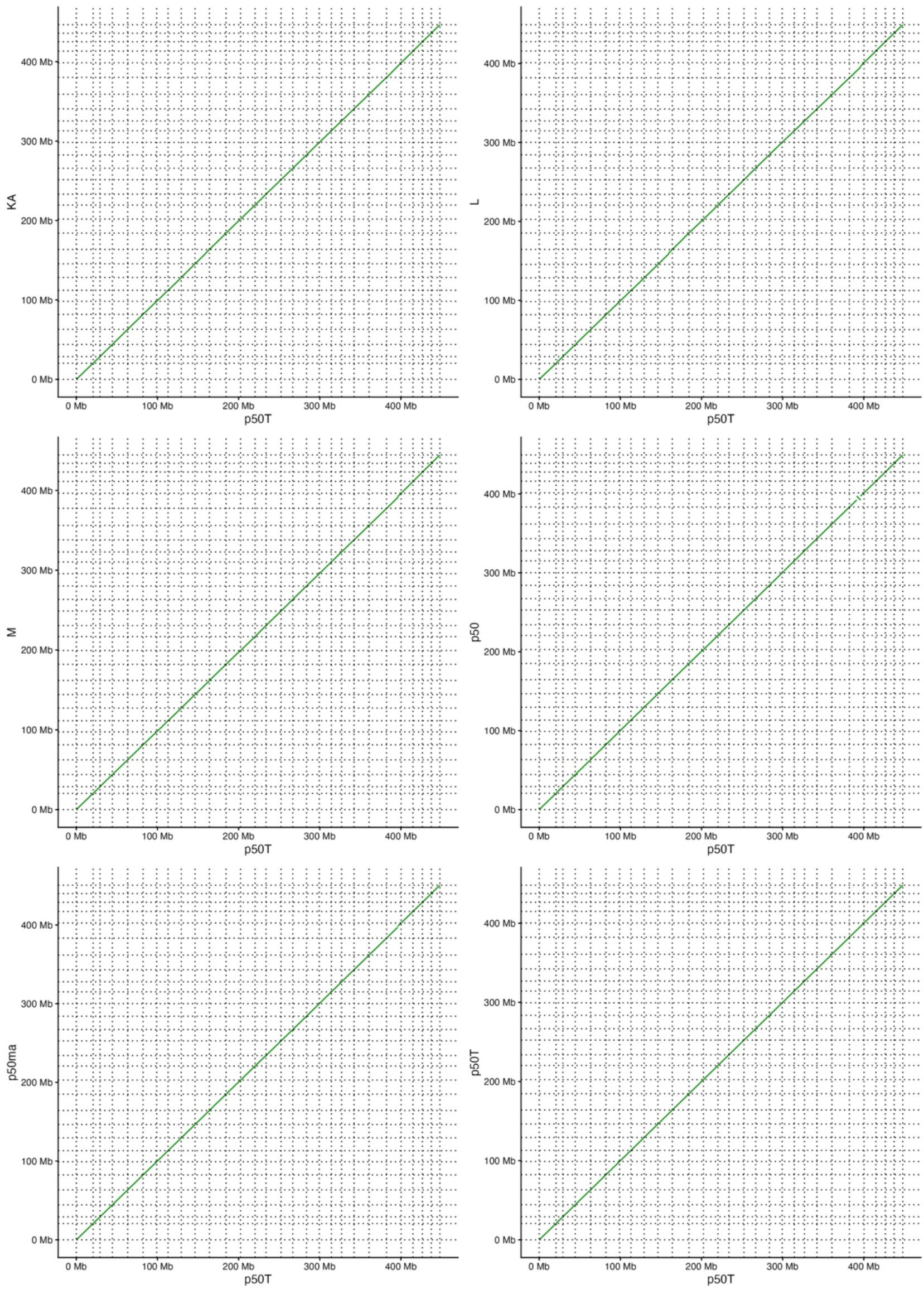
Whole-genome alignments among different *Bombyx* assemblies. The Z and autosomal chromosome scaffolds from various *Bombyx* genome assemblies were aligned using minimap2 (with default parameters) to a common reference (p50T) and visualized as dotplots. The last alignment (bottom right) is of p50T with itself, provided as a comparison to the other alignments between assemblies. Dotted lines delineate different chromosomes, which are arranged numerically from the origin (*e.g.,* Chr1/Z to Chr28).

## 2 Supplementary Tables

**Supplementary Table S1. GenBank accessions for *B. mori* and *B. mandarina* mitochondrial genomes used for principal component analysis.** Table provided as a separate Excel spreadsheet file.

**Supplementary Table S2.** GenBank Sequence Read Archive accessions for PacBio HiFi reads associated with each assembly analyzed directly in this manuscript.

| Assembly | Accession(s) |
| --- | --- |
| **Assembly** | DRR452101 |
| **p50ma** | SRR23490963 |
| **p50T<sup>a</sup>** | DRR092736, DRR092742-3, DRR092746-8, DRR092750-1, DRR092755, DRR092757, DRR092771-3 |

a) Multiple SRA entries were combined into a single data set; Ranges given by dashes indicate use of all data included in the range.

**Supplementary Table S3.**
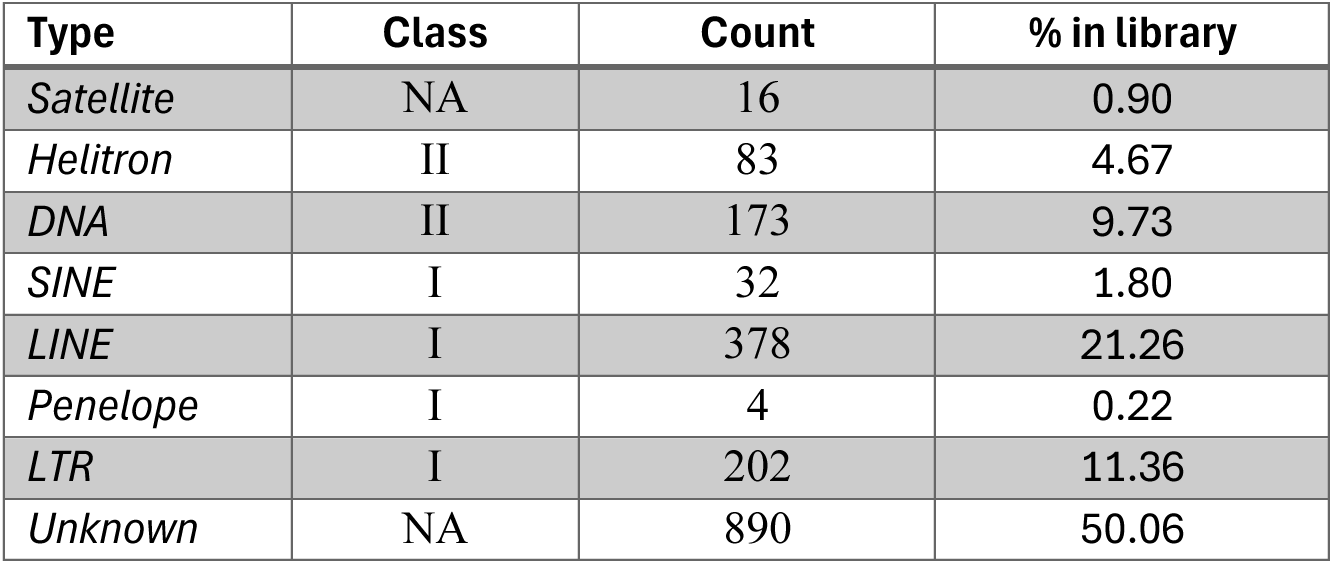
Composition of the *Bombyx* repeat library.

**Supplementary Table S4.** Percentage of repeat masked sequence for the W, Z, and autosomes in two *Bombyx* taxa.

| Taxon | Chromosome | % Repeat Masked |
| --- | --- | --- |
| *B. mandarina* | W | 95.8 |
|  | A | 48.1 |
|  | Z | 39.6 |
| *B. mori* | W | 95.4 |
|  | A | 50.4 |
|  | Z | 40.3 |

**Supplementary Table S5.** Percentage of repeat masked sequence for the W, Z, and autosomes in two *Bombyx* taxa, partitioned by type of repeat.

|  | *B. mori* |  |  | *B. mandarina* |  |  |
| --- | --- | --- | --- | --- | --- | --- |
| Repeat | W | A | Z | W | A | Z |
| DNA | 2.7 | 3.9 | 4.5 | 2.2 | 4.2 | 5.0 |
| Helitron | 0.6 | 3.6 | 3.2 | 0.7 | 3.6 | 3.2 |
| LINE | 41.0 | 20.5 | 15.8 | 39.5 | 19.4 | 15.1 |
| LTR | 39.9 | 6.6 | 4.8 | 42.4 | 5.4 | 4.2 |
| Penelope | 0.0 | 0.2 | 0.1 | 0.0 | 0.2 | 0.1 |
| SINE | 1.1 | 5.4 | 3.8 | 1.2 | 5.4 | 4.0 |
| Satellite | 4.5 | 0.1 | 0.1 | 1.9 | 0.0 | 0.0 |
| Unknown | 5.6 | 10.1 | 8.0 | 7.8 | 9.9 | 7.9 |

**Table S6.**
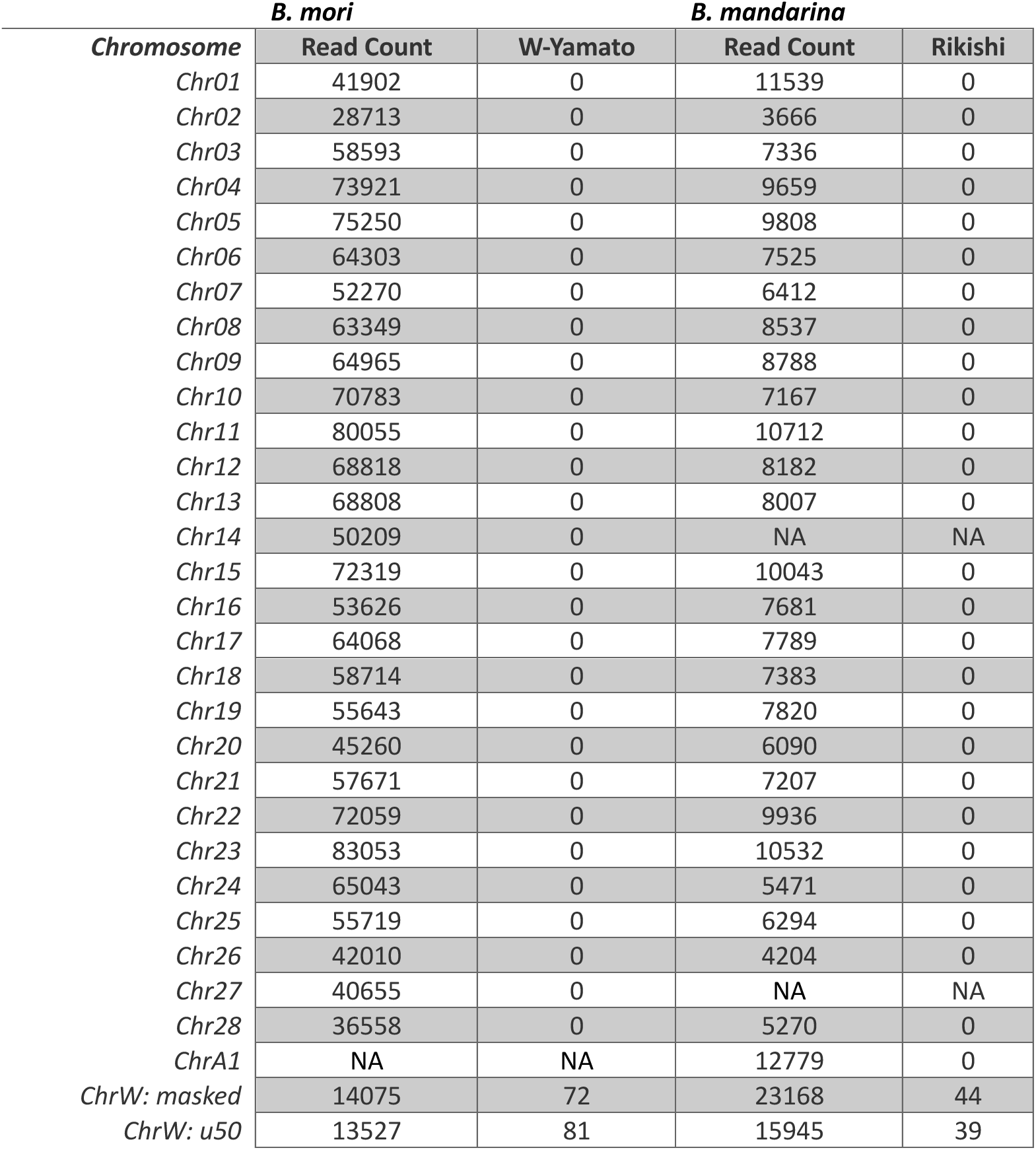
Counts of reads and the associated number of W-linked markers identified among reads assigned to each chromosome in PacBio sequencing data for *B. mori* and *B. mandarina*.

| *Chromosome* | *B. mori* |  | *B. mandarina* |  |
| --- | --- | --- | --- | --- |
|  | Read Count | W-Yamato | Read Count | Rikishi |
| *Chr01* | 41902 | 0 | 11539 | 0 |
| *Chr02* | 28713 | 0 | 3666 | 0 |
| *Chr03* | 58593 | 0 | 7336 | 0 |
| *Chr04* | 73921 | 0 | 9659 | 0 |
| *Chr05* | 75250 | 0 | 9808 | 0 |
| *Chr06* | 64303 | 0 | 7525 | 0 |
| *Chr07* | 52270 | 0 | 6412 | 0 |
| *Chr08* | 63349 | 0 | 8537 | 0 |
| *Chr09* | 64965 | 0 | 8788 | 0 |
| *Chr10* | 70783 | 0 | 7167 | 0 |
| *Chr11* | 80055 | 0 | 10712 | 0 |
| *Chr12* | 68818 | 0 | 8182 | 0 |
| *Chr13* | 68808 | 0 | 8007 | 0 |
| *Chr14* | 50209 | 0 | NA | NA |
| *Chr15* | 72319 | 0 | 10043 | 0 |
| *Chr16* | 53626 | 0 | 7681 | 0 |
| *Chr17* | 64068 | 0 | 7789 | 0 |
| *Chr18* | 58714 | 0 | 7383 | 0 |
| *Chr19* | 55643 | 0 | 7820 | 0 |
| *Chr20* | 45260 | 0 | 6090 | 0 |
| *Chr21* | 57671 | 0 | 7207 | 0 |
| *Chr22* | 72059 | 0 | 9936 | 0 |
| *Chr23* | 83053 | 0 | 10532 | 0 |
| *Chr24* | 65043 | 0 | 5471 | 0 |
| *Chr25* | 55719 | 0 | 6294 | 0 |
| *Chr26* | 42010 | 0 | 4204 | 0 |
| *Chr27* | 40655 | 0 | NA | NA |
| *Chr28* | 36558 | 0 | 5270 | 0 |
| *ChrA1* | NA | NA | 12779 | 0 |
| *ChrW: masked* | 14075 | 72 | 23168 | 44 |
| *ChrW: u50* | 13527 | 81 | 15945 | 39 |

## Notes

https://doi.org/10.5281/zenodo.21361476

### Competing Interest Statement

The authors have declared no competing interest.

